# One More Night: Investigating the Stability of Verbal Overnight Memory Consolidation and Sources of Variability

**DOI:** 10.64898/2026.11.23.767957

**Authors:** Nicholas Whelan, Aaron Lam, Clara Cher Yi Tan, David Ireland, Jurgen Fripp, Sharon L Naismith, Nathan Cross

## Abstract

Sleep-Dependent Memory Consolidation (SDMC) plays a vital role in facilitating the retention of newly learnt information. However, retesting in the literature is scarce and thus little is known regarding the daily stability of the SDMC process. The present study investigated whether overnight memory consolidation performance is stable (reflecting a trait) or variable (reflecting a state) across two nights using a novel word-pair smartphone application. Furthermore, the study examined the influence of subjective sleep quality and motivation on overnight consolidation performance, and whether these associations remained consistent across nights. Young healthy adults (*N* = 52) completed a validated word-pair task via a smartphone application over two nights and completed subjective sleep and motivation questionnaires. Results indicated overnight consolidation performance was not stable across nights, despite same-day and next-day recall exhibiting significant stability. Subjective sleep and motivation scores were not associated with performance, although these associations remained consistent across nights. However, worse PSQI scores were associated with lower next-day recall. After considering semantically related and unrelated word-pairs, both same-day and next-day recall performance exhibited significant stability for unrelated word-pairs only. These findings question the reliance on single-night SDMC protocols, and future research should examine modifiable factors that underlie SDMC variability across nights.

## INTRODUCTION

Memory formation involves both the initial encoding of information and its subsequent consolidation into long-term memory, which can later be evaluated through recall (Walker & Stickgold, 2004). While memories are initially formed and consolidated during wakefulness, a substantial body of research demonstrates that both circadian-aligned sleep and daytime napping play a vital role in their subsequent strengthening and reorganisation, a process known as sleep-dependent memory consolidation (SDMC) (Stickgold, 2005).

The word-pair recall task is a prominently used paradigm in the literature for investigating verbal SDMC (Rasch & Born, 2013). Typically, participants are sequentially presented with a series of word-pairs during an initial learning phase. Following all items, they are then tested by being presented with the first word as a cue, and must recall the second word of each pair. To assess SDMC, testing is performed both before and after sleep. The standard metric for SDMC is calculated by dividing (Holz et al., 2012; Lam et al., 2021; Teh et al., 2023; Wilhelm et al., 2011), or subtracting (Backhaus et al., 2008; Feld et al., 2016; Gais & Born, 2004; Lo et al., 2014; van Schalkwijk et al., 2019; van Sluijs et al., 2020; Wilhelm et al., 2008) the morning performance by the pre-sleep performance to create a normalised overnight retention score that captures the consolidation process between short-term (e.g. 30 min) and longer-term (e.g. hours) storage (Lam et al., 2021; van Sluijs et al., 2020). Figure 1 illustrates an example procedure.

**Figure 1.**
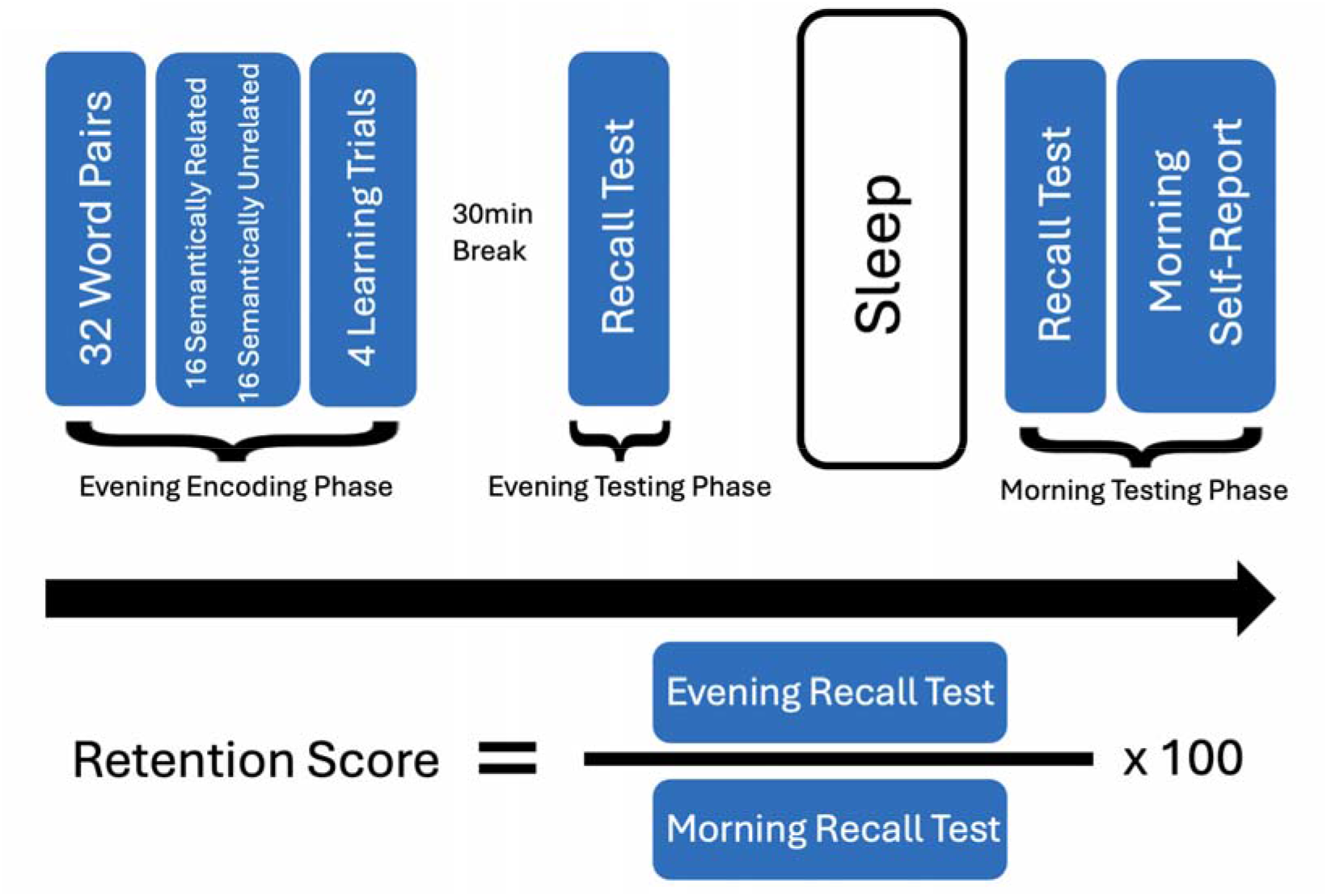
Word-Pair Task Paradigm. *Note.* Diagram adapted from Lam et al. (2021) showing word-pair task procedure. Retention Score shows calculation of a normalised overnight retention score for SDMC.

Results generally show that sleep has a beneficial influence on the number of word-pairs recalled (Gais & Born, 2004; Payne et al., 2012; Tucker & Fishbein, 2008; Wilhelm et al., 2008; Wilson et al., 2012). Importantly, word-pair semantics modulate the beneficial influence. Semantically unrelated word-pairs (e.g. “black” – “standing”) show minimal SDMC improvement compared to semantically related pairs (e.g. “mother” – “daughter”) (Gais & Born, 2004; Payne et al., 2012). This difference is likely because related word-pairs can leverage pre-formed hippocampal associations (Stickgold, 2004).

Although SDMC testing paradigms are well established, a fundamental assumption has remained untested: that recall performance after one night is representative of an individual’s typical performance on the task. A large proportion of studies assess baseline levels of SDMC on one occasion, with no retesting across nights (Rasch & Born, 2013). Without retesting, it becomes difficult to determine if performance reflects state-dependent daily factors, or instead reflects stable, trait-dependent factors within the individual that are resistant to fluctuations (Nemeth et al., 2024). Importantly, sleep-manipulation studies using targeted memory reactivation (Schreiner & Rasch, 2015), closed-loop auditory stimulation (Ngo et al., 2013), and all-night rocking (Perrault et al., 2019) have also relied on the assumption that performance during non-experimental or baseline nights provides a stable reference against which manipulation-related changes can be evaluated. Indeed, findings from these manipulation studies demonstrate that SDMC may be sensitive to experimental influences and can vary meaningfully across nights, however evidence is still mixed. While some studies report benefit of experimental manipulation for memory consolidation (Marshall et al., 2006; Ngo et al., 2013; Schreiner & Rasch, 2015; Papalambros et al., 2017; Perrault et al., 2019; Westerberg et al., 2015), others have reported little to no benefit (Bueno-Lopez et al., 2019; Diep et al., 2020; Eggert et al., 2013; Harrington et al., 2021; Henin et al., 2019; Koo et al., 2018; Omlin et al., 2018; Paßmann et al., 2016; Sahlem et al., 2015; Schneider et al., 2020; van Sluijs et al., 2020; Wilhelm et al., 2020). Furthermore, a meta-analysis by Harlow et al. (2023) highlights a yearly decline in effect size for acoustic stimulation studies addressing memory retention. Thus, mixed findings across manipulation studies may partly reflect the use of a single baseline assessment for a process that fluctuates within individuals, such that the particular night on which baseline performance is measured may influence the apparent effect of the experimental manipulation.

Furthermore, these findings do not directly establish whether individual differences in SDMC are stable over time. Trait-like abilities have already been established in other cognitive domains through retesting. General intelligence has been shown to be stable across timespans ranging from weeks to years (Breit et al., 2024; Danner et al., 2011; Larsen et al., 2008). Same-day memory tasks, such as the Rey Auditory Verbal Learning Test (RAVLT) (Rey, 1958) and the Wechsler Memory Scale Verbal Paired Associates (VPA) (Wechsler, 2009) have both shown moderate to high test-retest validity (Lee et al., 2023; Magalhaes et al., 2012; Rezvanfard et al., 2011). Importantly, these tasks use procedures and word-pairs comparable to SDMC evening testing. Therefore, it is plausible that sleep-dependent memory exhibits similar stability to same-day learning rates and recall. However, the lack of SDMC trait analyses means little is known about its stability.

If SDMC is not completely trait-like, then night-to-night variation may obscure associations with metrics such as sleep quality. Although objective sleep features such as NREM sleep spindles have been associated with memory consolidation, findings are mixed (Ujma, 2024), suggesting that SDMC variability may obscure any relationships – especially given that sleep microarchitecture features are known to be highly stable and trait-like. Repeated polysomnography is costly and difficult to implement (Kumral et al., 2023; Schabus et al., 2004; Samson, 2021), however, subjective sleep measures offer an easy, scalable alternative. Identifying an individual’s baseline SDMC by calculating an average SDMC performance score across nights may reduce noise and strengthen associations with subjective reports of sleep quality, which could have potential clinical utility.

Accordingly, the present study used a smartphone application (Ireland et al., 2024, Tan et al., 2026) to examine the stability of SDMC across multiple nights, benchmarked against pre-sleep learning and recall. We also examined whether subjective sleep quality predicts SDMC within and across nights, using home-based assessments that enable scalable measurement under ecologically valid conditions. It was hypothesised that SDMC would show moderate-to-strong stability across nights, comparable to the stability of pre-sleep learning and recall, while better subjective sleep quality would be associated with better SDMC across nights. Furthermore, it was hypothesised that stability and associations with sleep quality and motivation would be greater for semantically related word pairs than unrelated word pairs.

## METHODS

### Participants

Undergraduate psychology students were recruited from the University of Sydney’s SONA Psych pool, participating for course credit. All procedures were approved by the University of Sydney Human Research Ethics Committee (2025/299).

Participants must have been free of any major neurological conditions (e.g. epilepsy, stroke, brain injury), sleep disorders (e.g. obstructive sleep apnoea, insomnia, narcolepsy), or cognitive impairment (MMSE score < 25, MoCA score < 26). Participants were also expected to maintain a regular sleep schedule with bedtimes between 9pm – 1am and rise times between 6am – 10am.

### Materials

#### Subjective Sleep Quality

The Pittsburgh Sleep Quality Index (PSQI) (Buysse et al., 1989) is a 19-item self-report questionnaire assessing sleep quality, duration, latency, disturbances, medication, daytime dysfunction, and habitual sleep efficiency from the past month. Total scores range from 0-21. Higher scores indicate worse sleep quality. Additionally, participants completed a one-item daily morning self-report rating of their sleep quality the night prior. Scores range from 0-10, with higher scores indicating greater sleep quality.

#### Motivation

Motivation may influence encoding and the preferential consolidation of salient memories, potentially confounding repeated assessments of SDMC (Duan et al., 2020; Rijn et al., 2017; Studte et al., 2017; Wang et al., 2024; Wilhelm et al., 2011). Therefore, we collected subjective measures of motivation and apathy to control for differences across participants. The Apathy-Motivation Index (AMI) (Ang et al., 2017) is an 18-item questionnaire measuring apathy and motivation in healthy individuals based on the past two weeks. AMI subscales relate to behavioural activation, social motivation, and emotional sensitivity. Total scores range between 0-72, with higher scores indicating greater apathy and lack of motivation. Participants also completed a one-item daily morning self-report rating of their motivation the day prior. Scores range from 0-10, with higher scores indicating greater motivation.

#### Depression

Depression was collected for demographic purposes using the Patient Health Questionnaire-9 (PHQ-9) (Kroenke et al., 1999), a nine-item questionnaire measuring severity of depression. Scores range between 0-27 with higher scores indicating greater depressive symptoms.

#### Cognitive Screening

Participants were screened for cognitive impairment using the Montreal Cognitive Assessment (MoCA) (Nasreddine et al., 2005) or Mini-Mental State Examination (MMSE) (Folstein et al., 1975). Scores range from 0-30 for both assessments, lower scores indicate worse cognitive functioning, with scores < 25 for the MMSE and < 26 for the MoCA indicating cognitive impairment.

#### SleepMemories Application

To assess SDMC, data was collected using the SleepMemories smartphone application, which administered a word-pair task adapted from Lam et al. (2021). The task included 32 word-pairs, 16 semantically related (e.g. “mother” – “daughter”), and 16 semantically unrelated (e.g. “black” – “standing”). The application presents all participants with the same word-pair list on the first night of testing.

### Design

This study employed a repeated-measures design where young adults performed a word-pairs task before and after sleep across two nights. Participants were screened for cognitive impairment using either the MoCA or MMSE before providing informed consent. Participants completed the PHQ-9, AMI, and PSQI before installing the SleepMemories smartphone application. Participants were informed to complete tasks on the application across two nights, with successful completion resulting in a draw entry to win one of four $50 cash prizes. Each morning, participants completed a self-report on their motivation and sleep quality the night prior.

### Experimental Procedure

#### Setup

After installing and opening the SleepMemories application, participants answered demographic questions and entered their expected sleep and wake-up times.

#### Evening Encoding Phase

Approximately 3 hours prior to their nominated sleep time, participants received a notification to complete the evening (pre-sleep) learning trials. Participants completed four learning trials, in which they were randomly presented 32 word-pairs, one at a time, at the rate of one word-pair every 10 seconds. After each learning trial, participants were presented with the first word of each word-pair and required to enter the corresponding word within 15 seconds. See Figure 2 for example of word-pair learning trial tasks.

**Figure 2.**
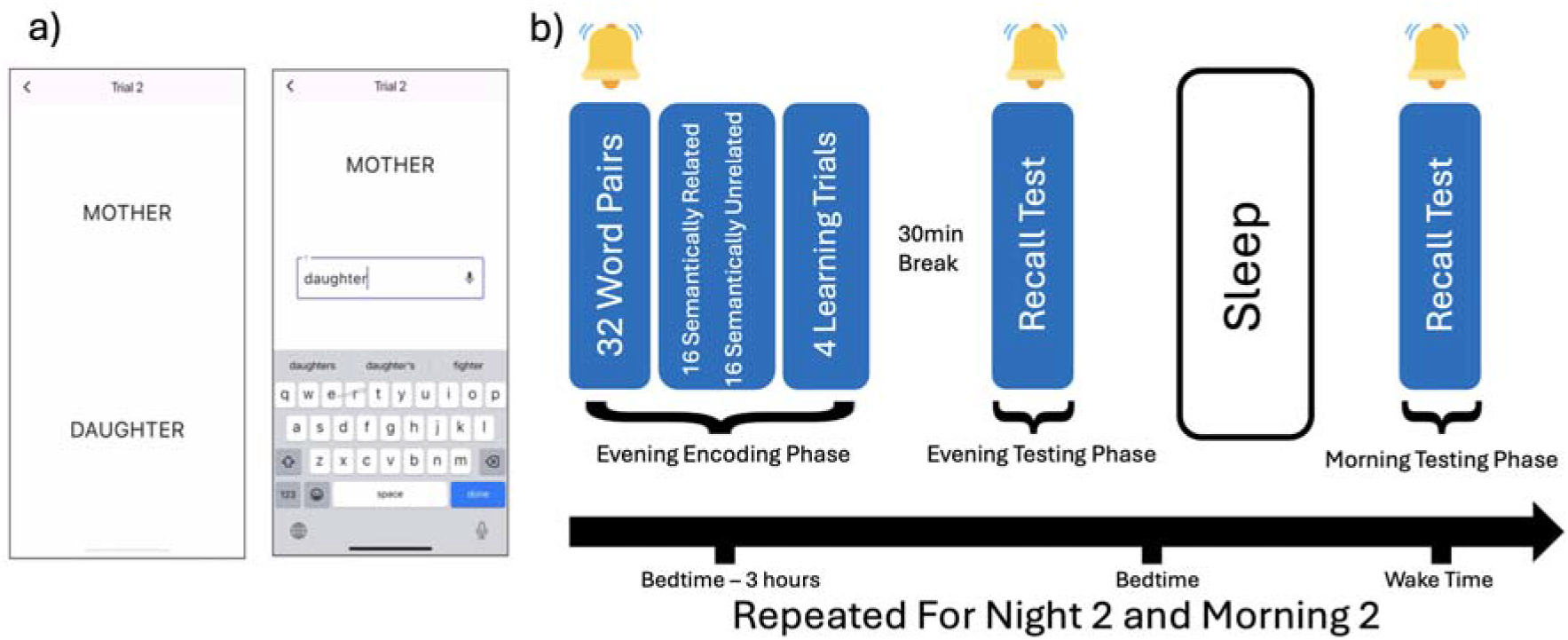
Example of Word-Pair Learning Trial Task. *Note*. a) Word-pair learning and recall example. The participant is shown the first word and prompted to enter the corresponding word. b) Word-pair task procedure. The same procedure was repeated across two nights to assess stability, with separate wordlists for each night.

#### Evening Testing Phase

Following encoding, participants had a thirty-minute break before they were again notified to complete the evening (pre-sleep) recall test. Participants were randomly presented with the first word of each word-pair and required to enter the corresponding word.

#### Morning Testing Phase

At the nominated wake-up time, participants received a notification to complete the morning recall test, which followed the same format as the evening recall test. To test the stability of SDMC, participants repeated the same process on a different evening and morning (within a week), learning a separate, randomly assigned word-pair list (one of 3 alternatives) (Figure 2), along with daily morning self-report ratings for sleep and motivation.

### Analysis Plan

#### Primary and Exploratory Outcomes

The primary outcome for SDMC was a normalised overnight retention score calculated as the percentage of words recalled in the morning, compared to pre-sleep recall (e.g. (Morning Recall/Pre-Sleep Recall) x 100, see Figure 1). Baseline measures included pre-sleep recall and learning rate (slope of learning trials), serving as stability benchmarks for same-day memory. Morning recall performance was also included for analyses.

#### Statistical Analysis

All analyses were conducted in R, Version 4.5.1 (R Core Team, 2025), using the packages tidyverse (Wickham et al., 2019), psych (Revelle, 2025), and rstatix (Kassambara, 2023). Normality and homogeneity of variance were checked using the Shapiro-Wilk test and Levene’s test, respectively. Given the focus on within-subject stability and the potential for anomalous observations during unsupervised, at-home assessment, multivariate outliers were identified and removed using Mahalanobis distance (Mahalanobis, 1936/2018). Participants with minor missing data were retained, with data excluded from relevant analyses using pairwise deletion. Stability of SDMC was evaluated using Spearman’s correlations, intraclass correlation coefficients (ICC), and paired Wilcoxon tests, assessing relative stability and night-to-night differences, respectively. The relationship between these measures and PSQI and AMI scores was analysed using Spearman correlations. Correlations were performed for each day separately, and their average. Fishers r to z were conducted to test differences in associations between nights. Daily morning self-reports for sleep quality and motivation were included for exploratory analyses of night-specific influences and correlated with corresponding night SDMC measures. Paired Wilcoxon signed-rank tests were run to check differences in learning and testing start times between nights.

## RESULTS

### Participants

A total of 87 participants were recruited for the study. Data from 35 participants were excluded due to app-related software issues (n = 30) or missing/invalid task completion (n = 5). Multivariate outliers were identified using Mahalanobis distance for Night 1 and Night 2 recall scores, learning rate, and overnight retention. Participants that exceeded the chi-square value (χ² = 9.21, df = 2, p = .01) were flagged as outliers. A total of five outliers were removed before analyses, three based on overnight retention scores and two based on morning recall. One participant was flagged as an outlier across multiple variables. The final analytic sample consisted of 52 participants. Participants were aged 18–33 years (M = 20.6, SD = 2.4), and 81% were female (n = 42). All participants performed within the intact cognitive range on either the MMSE (range = 29–30, n = 34) or MoCA (range = 26–29, n = 18). PHQ-9 scores ranged from 1–21 (M = 5.0, SD = 4.5), and PSQI scores ranged from 3–14 (M = 6.0, SD = 2.6). Among participants who completed the AMI, scores ranged from 8–39 (M = 25.2, SD = 8.0).

### Night-to-Night Stability of Memory Performance

Results indicated significant positive night-to-night correlations for Pre-Sleep Learning Rate, *r_s_*(49) = .65, *p* < .001 (Figure 3), Pre-Sleep Recall Performance, *r_s_*(42) = .51, *p* < .001, and Morning Recall Performance, *r_s_*(44) = .58, *p* < .001. In contrast, there was no significant night-to-night correlation for the primary normalised overnight retention score, *r_s_*(38) = .11, *p* = .489.

**Figure 3.**
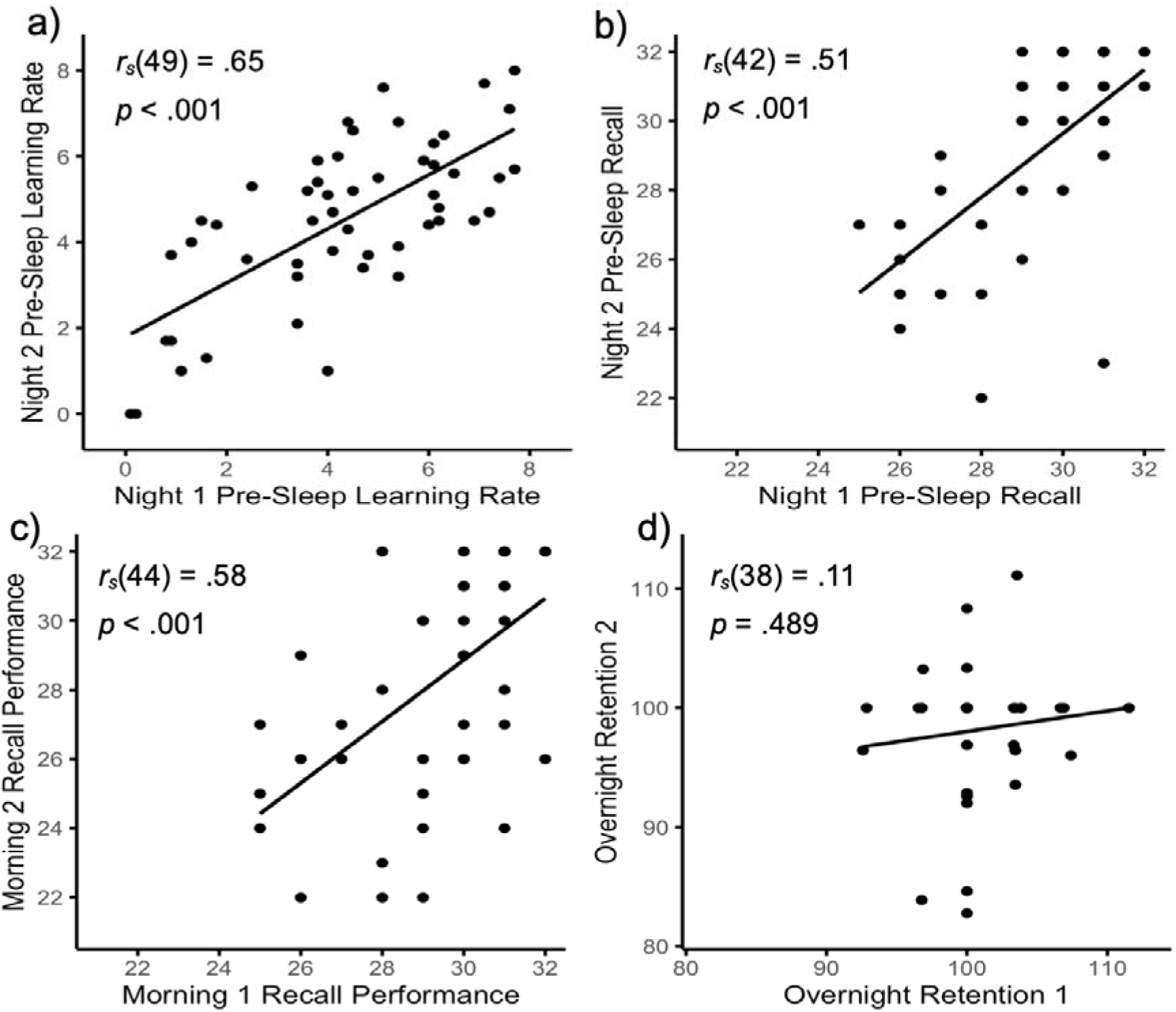
Scatterplots of Day 1 vs Day 2 Memory Measures. *Note*. Scatterplots of a) Night 1 Pre-Sleep Learning Rate vs Night 2 Pre-Sleep Learning Rate, b) Night 1 Pre-Sleep Recall vs Night 2 Pre-Sleep Recall, c) Morning 1 Recall vs Morning 2 Recall, d) Night 1 Overnight Retention Score vs Night 2 Overnight Retention Score.

Intraclass correlation coefficients (ICCs) further supported night-to-night consistency in task performance measures. Consistency was highest for pre-sleep learning rate, ICC = .82, 95% CI [.69, .90], followed by morning recall performance, ICC = .65, 95% CI [.38, .80], and pre-sleep recall performance, ICC = .59, 95% CI [.28, .76]. In contrast, the overnight retention score showed weaker evidence of night-to-night consistency, ICC = .35, 95% CI [−.13, .63].

Paired Wilcoxon signed-rank tests indicated no significant night-to-night differences in pre-sleep recall performance, or learning rate. However, both overnight retention and morning recall performance were significantly higher on Night 1 than Night 2, with a moderate effect size (Table 1). Subjective sleep quality and motivation ratings also did not differ significantly between nights.

**Table 1.**
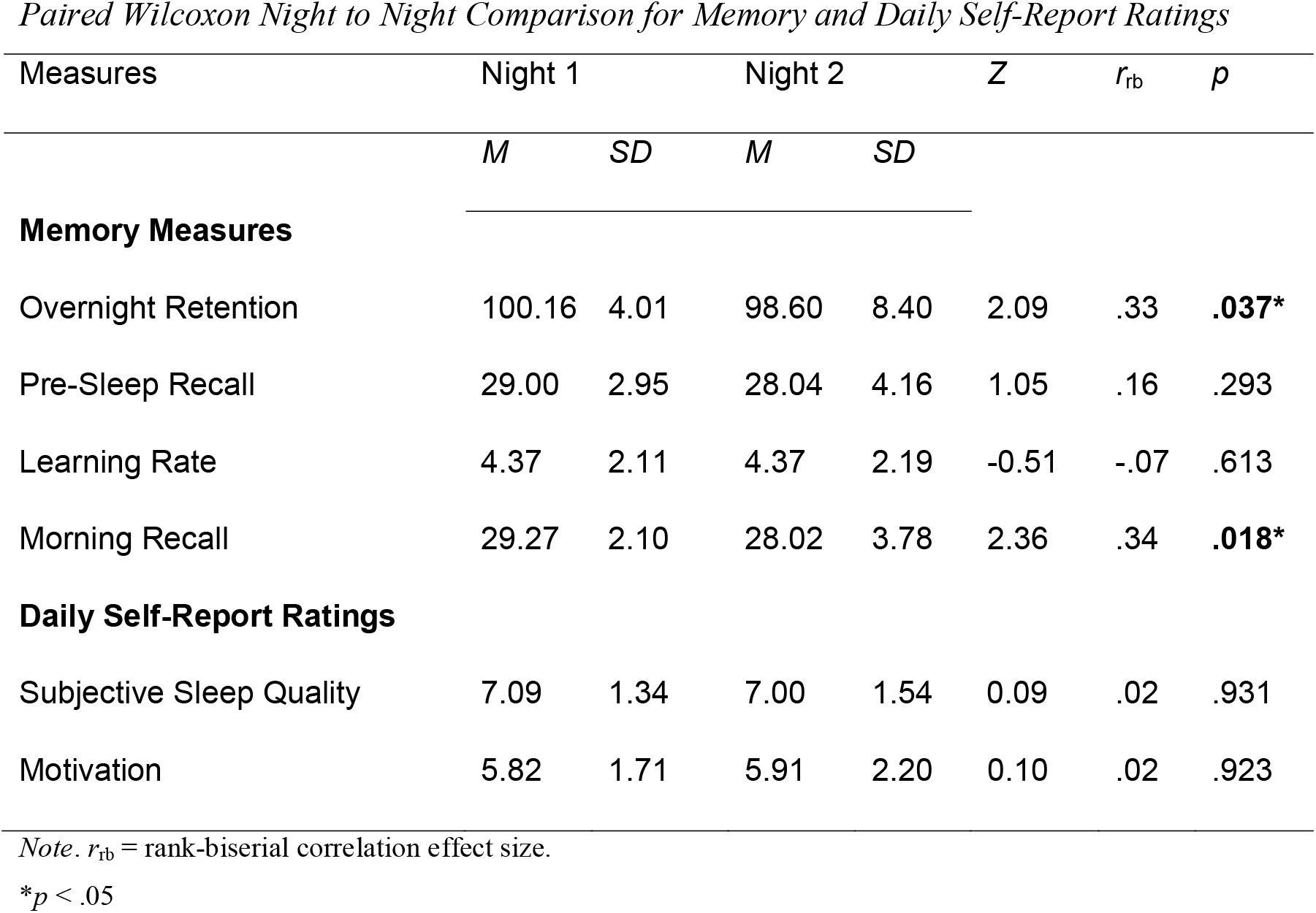
Paired Wilcoxon Night to Night Comparison for Memory and Daily Self-Report Ratings.

Bland–Altman analyses indicated small average night-to-night biases across memory measures, suggesting minimal systematic change between Night 1 and Night 2. Mean bias was -2.93 for overnight retention, -1.07 for morning recall, -0.84 for pre-sleep recall, and 0.19 for pre-sleep learning rate. However, the limits of agreement indicated considerable individual variability, particularly for overnight retention, where scores varied from −15.63 to 9.79 across nights. Limits of agreement were narrower for morning recall [−6.33, 4.20], pre-sleep recall [−7.28, 5.60], and pre-sleep learning rate [−2.91, 3.28]. (Figure 4).

**Figure 4.**
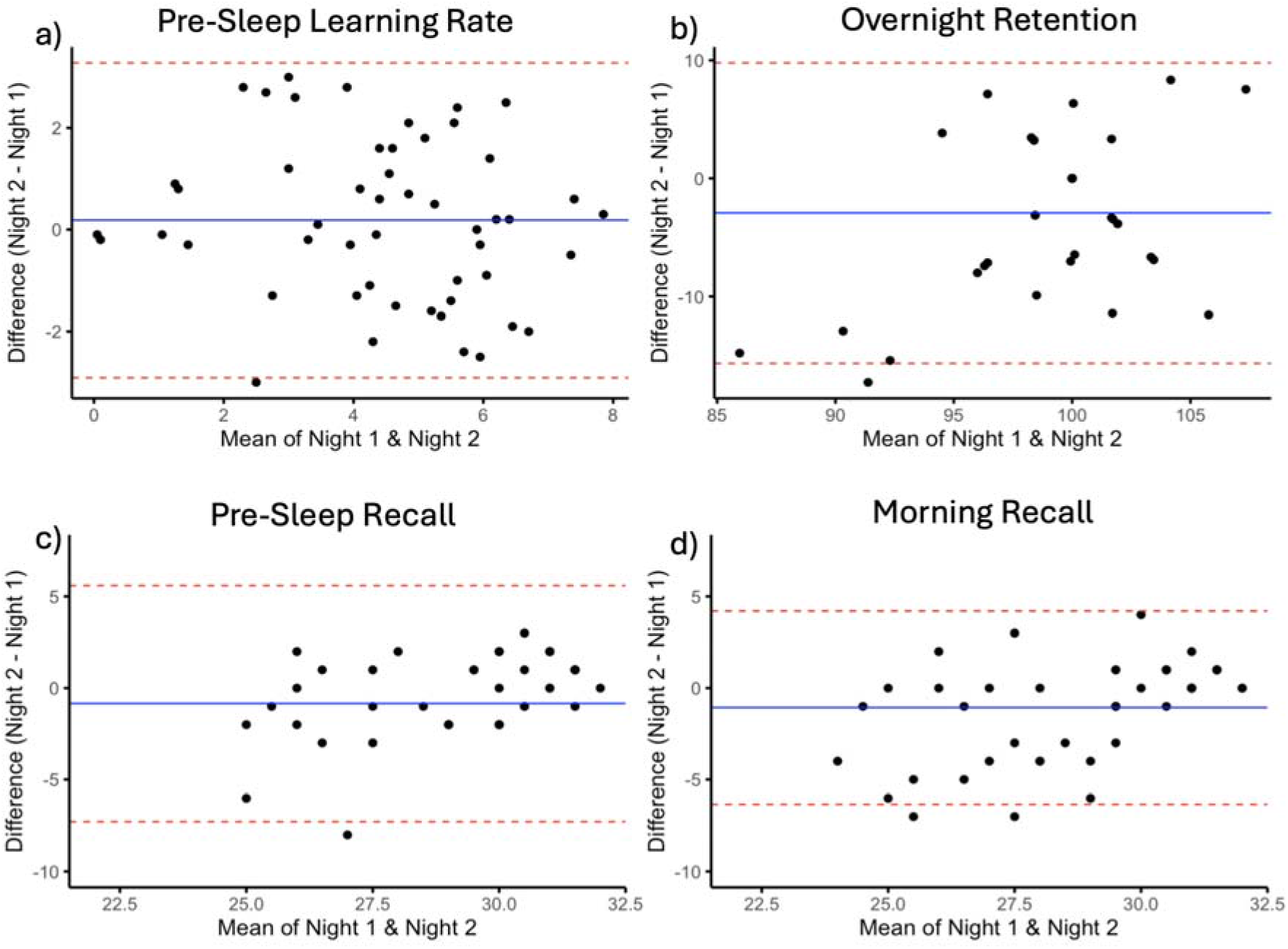
Bland-Altman Plots of Night 1 vs Night 2 Measures. *Note*. Bland-Altman Plots of a) Night 1 Pre-Sleep Learning Rate vs Night 2 Pre-Sleep Learning Rate, b) Overnight Retention Score 1 vs Overnight Retention Score 2, c) Night 1 Pre-Sleep Recall vs Night 2 Pre-Sleep Recall, d) Morning 1 Recall vs Morning 2 Recall. The x-axis represents the mean performance across Night 1 and Night 2, and the y-axis represents the difference between nights (Night 2 − Night 1). Solid lines indicate the mean difference (bias), and dashed lines indicate the 95% limits of agreement.

### Timing Checks

There were no significant differences in start times for Pre-Sleep Learning, Pre-Sleep Testing, or Morning Testing. However, the gap between Pre-Sleep Learning and Testing was found to be significantly different in Night 2 despite identical means. Pre-Sleep Learning started at ∼20:57 (±1h 34min) on Night 1 and ∼20:44 (±3h 12min) on Night 2. Pre-Sleep Testing commenced at ∼22:11 (±1h 56min) on Night 1 and ∼22:23 (±1h 32min) on Night 2. Morning Testing started at ∼9:14 (±3h 9min) for the first Morning and ∼9:38 (±2h 45min) the second. The gap between pre-sleep learning and pre-sleep testing for Night 1 was on average 68.39 minutes (±90.80 min) and 68.39 minutes (±31.13 min) for Night 2 (Table 2).

**Table 2.** Paired Wilcoxon Night to Night Comparison for Test Times.

| Measures | Night 1 |  | Night 2 |  | *Z* | *r*<sub>rb</sub> | *p* |
|---|---|---|---|---|---|---|---|
|  | *M* | *SD* | *M* | *SD* |  |  |  |
| **Test Times (time)** |  |  |  |  |  |  |  |
| Pre-Sleep Learning to Pre-Sleep Test Gap | 1:08:23 | 90.80 | 1:08:23 | 31.13 | -2.91 | -.44 | **.004\*\*** |
| Pre-Sleep Learning | 20:57:16 | 94.11 | 20:43:54 | 191.96 | -0.70 | -.10 | .486 |
| Pre-Sleep Testing | 22:10:50 | 115.82 | 22:23:13 | 91.74 | -1.38 | -.21 | .167 |
| Morning Testing | 9:14:16 | 188.86 | 9:38:23 | 164.78 | -1.67 | -.25 | .095 |

*Note*. *r*<sub>rb</sub> = rank-biserial correlation effect size. Test time means shown as HH:MM:SS. SD represented as minutes.
\*\**p* < .01

No significant correlation was found between the difference in morning test times and difference in overnight retention, (*r_s_*(36) = .14, *p* = .415), or the difference in morning recall performance (*r_s_*(42) = -.09, *p* = .559). Furthermore, no significant correlation was found between the difference in pre-sleep learning-test gap and difference in overnight retention, (*r_s_*(36) = .04, *p* = .833), or morning recall performance (*r_s_*(39) = -.13, *p* = .430).

### Relationship between SDMC and Sleep Quality

No significant correlation was found with the PSQI global score and overnight retention on Night 1, (*r_s_*(43) = .16, *p* = .305), Night 2, (*r_s_*(38) = .06, *p* = .727), or their average, (*r_s_*(37) = .12, *p* = .462). A Fisher’s r-to-z revealed no significant difference between correlation coefficients for Night 1 and Night 2, (*Z* = 0.45, *p* = .653).

For morning recall performance, a significant correlation was found with PSQI for Morning 1, (*r_s_*(46) = -.32, *p* = .026, *q* = .030), Morning 2, (*r_s_*(44) = -.32, *p* = .030, *q* = .030), and their average, (*r_s_*(44) = -.38, *p* = .009, *q* = .028), these also survived correction for multiple comparisons (Figure 5). A Fisher’s r to z revealed no significant difference between correlation coefficients for Morning 1 and Morning 2, (*Z* = -0.005, *p* = .996). Furthermore, no significant correlation was found between variability in retention (Morning 2 – Morning 1) and PSQI (*r_s_*(37) = -.10, *p* = .559). For pre-sleep recall, a significant correlation was found with PSQI for Night 1, (*r_s_*(45) = -.40, *p* = .005, *q* = .016), and the average, (*r_s_*(40) = -.31, *p* = .044, *q* = .070), but not Night 2, (*r_s_*(41) = -.28, *p* = .070, *q* = .070). Only Night 1 remained significant after Hochberg correction. A Fisher’s r-to-z revealed no significant difference between correlation coefficients for Night 1 and Night 2, (*Z* = -0.63, *p* = .529).

**Figure 5.**
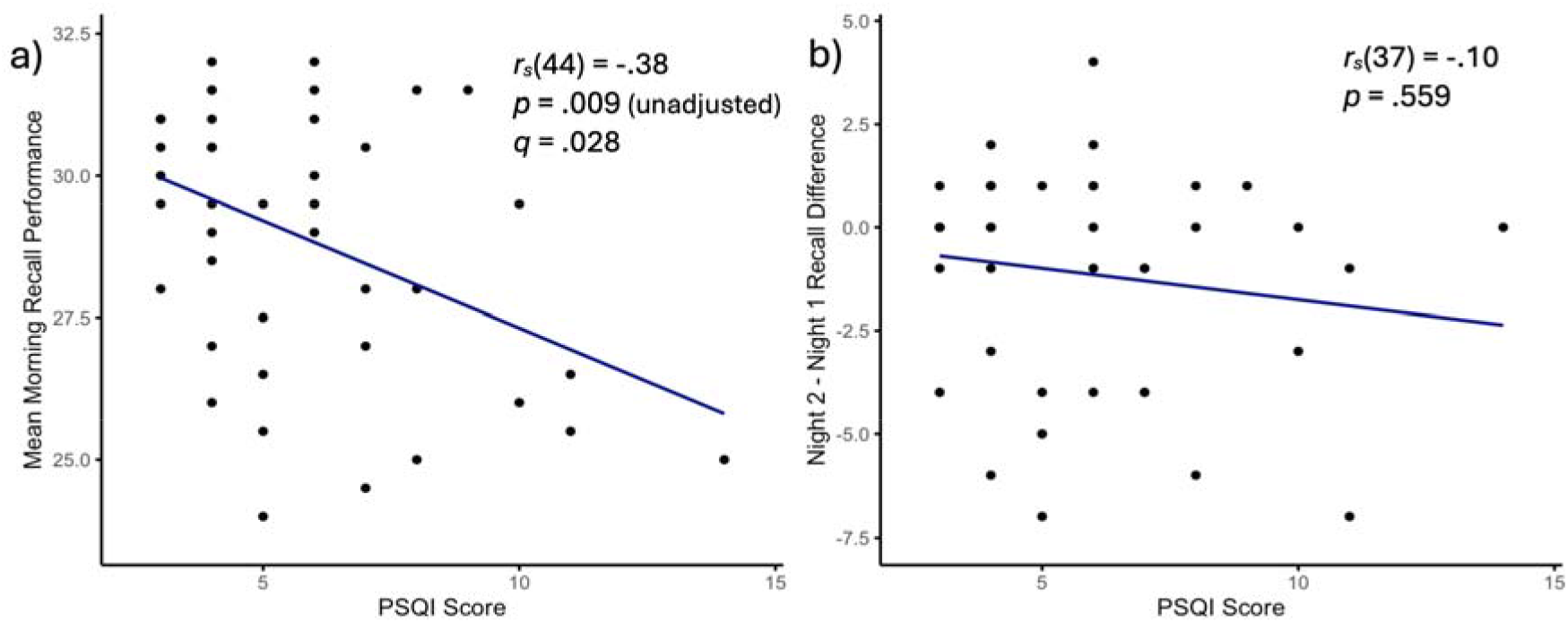
Scatterplots of Performance Measures vs PSQI Score. *Note*. Scatterplots of a) Average Morning Recall vs PSQI, b) Overnight Retention Variability (Night 2 – Night 1 Retention) vs PSQI.

Exploratory analyses for daily morning subjective sleep ratings (*n* = 22) showed no significant correlation between SDMC measures and reported sleep quality for each corresponding night, *p* > .05. Daily subjective sleep ratings were not significantly correlated across nights, *r_s_*(20) = .17, *p* = .445, and there was no significant nightly difference (Table 1).

### Relationship between SDMC and Motivation

For overnight retention, no significant correlation was found with the AMI score for Night 1, *r_s_*(29) = -.01, *p* = .961, Night 2, *r_s_*(28) = -.07, *p* = .706, or their average, *r_s_*(27) = -.13, *p* = .500. A Fisher’s r to z revealed no significant difference between correlation coefficients for Night 1 and Night 2, *Z* = 0.23, *p* = .815. No significant correlations were found between AMI and exploratory measures, *p* > .05. Further correlations were run between Behavioural Activation subscales of the AMI and memory measures. Results showed no significant correlations with any measures, *p* > .05. Fisher’s r to z revealed no significant difference in correlation coefficients between nights for any measure, *p* > .05. Exploratory analyses for daily morning motivation ratings (*n* = 22) showed no significant correlation between SDMC measures and motivation for each corresponding night, *p* > .05. Daily motivation self-reports did not differ between nights and were significantly correlated across nights, *r_s_*(20) = .67, *p* < .001.

#### Word pair semantics

For semantically related word-pairs, results indicated significant positive night-to-night correlations for pre-sleep learning rate, *r_s_*(49) = .523, *p* < .001. In contrast, there was no significant night-to-night correlation for the primary overnight retention score outcome, *r_s_*(37) = -.25, *p* = .128, morning recall performance, *r_s_*(44) = -.05, *p* = .764, or pre-sleep recall performance, *r_s_*(42) = -.02, *p* = .909. ICCs supported this pattern of consistency in task performance measures. Consistency was high for pre-sleep learning rate, ICC = .70, 95% CI [.47, .83]. In contrast, pre-sleep recall had low consistency, ICC = .03, 95% CI [-.69, .44], and all other ICC measures for related word pairs had an ICC of 0.00, 95% CI [-.74, .43].

Bland-Altman analyses for related word pairs revealed minimal mean bias across memory measures. Mean bias was -1.62 for overnight retention, -0.37 for morning recall, -0.09 for pre sleep recall, and -0.08 for pre-sleep learning rate. Limits of agreement were largest for overnight retention [-18.59,15.36], followed by pre-sleep recall [-2.82, 2.64], morning recall [-2.83, 2.09], and pre-sleep learning rate [-1.92, 1.75].

For semantically unrelated word-pairs, results also indicated significant positive night-to-night correlations for pre-sleep learning rate, *r_s_*(49) = .57, *p* < .001. However, there were also significant correlations for morning recall performance, *r_s_*(45) = .62, *p* < .001 and pre-sleep recall performance, *r_s_*(44) = .51, *p* < .001. There was no significant night-to-night correlation for overnight retention score, *r_s_*(37) = .18, *p* = .270. ICCs supported night-to-night consistency in task performance measures. Consistency was highest for pre-sleep learning rate, ICC = .78, 95% CI [.62, .87], followed by morning recall performance, ICC = .72, 95% CI [.51, .84], and pre-sleep recall performance, ICC = .67, 95% CI [.43, .81]. Overnight retention score showed weaker evidence of night-to-night consistency, ICC = .39, 95% CI [−.07, .65].

Bland-Altman analyses showed that for unrelated word pairs, mean bias was -3.69 for overnight retention, -0.91 for pre-sleep recall, -0.87 for morning recall, and 0.27 for pre-sleep learning rate. Overall, limits of agreement were wider than those for related word pairs. Again, overnight retention had the widest limits of agreement [-24.45, 18.07], followed by pre-sleep recall [-6.15, 4.33], morning recall [-5.11, 3.37], and pre-sleep learning rate [-1.94, 2.48].

Paired Wilcoxon signed-rank tests indicated no significant night-to-night differences in pre-sleep recall performance, learning rate, or overnight retention for both related and unrelated word-pairs. However, morning recall performance was significantly higher on Night 1 than Night 2 for both related and unrelated word-pairs, with a moderate effect size (Table 3).

**Table 3.** Paired Wilcoxon Night to Night Comparison for Memory Measures (Semantically Related and Unrelated Word-Pairs)

| Measures | Night 1 |  | Night 2 |  | *Z* | *r*<sub>rb</sub> | *p* |
|---|---|---|---|---|---|---|---|
|  | *M* | *SD* | *M* | *SD* |  |  |  |
| **Semantically Related Pairs** |  |  |  |  |  |  |  |
| Overnight Retention | 101.49 | 5.03 | 99.65 | 9.15 | 0.94 | 0.147 | .345 |
| Pre-Sleep Recall | 15.29 | 1.06 | 15.09 | 1.60 | 0.28 | 0.041 | .782 |
| Learning Rate | 1.50 | 0.88 | 1.30 | 1.28 | 0.90 | 0.125 | .368 |
| Morning Recall | 15.43 | 1.06 | 15.00 | 1.55 | 2.20 | 0.314 | **.028\*** |
| **Semantically Unrelated Pairs** |  |  |  |  |  |  |  |
| Overnight Retention | 99.93 | 10.82 | 97.58 | 12.74 | 1.14 | 0.179 | .253 |
| Pre-Sleep Recall | 13.71 | 2.43 | 12.96 | 3.07 | 1.77 | 0.261 | .077 |
| Learning Rate | 2.87 | 1.42 | 3.07 | 1.26 | 1.27 | 0.176 | .204 |
| Morning Recall | 13.53 | 2.27 | 12.49 | 3.37 | 2.68 | 0.382 | **.007\*\*** |

*Note*. *r*<sub>rb</sub> = rank-biserial correlation effect size.
\**p* < .05
\*\**p* < .01

For both semantically related and unrelated overnight retention scores, no significant correlation was found with PSQI for Night 1, Night 2, or their average, *p* > .05. For semantically related morning recall scores, a significant correlation with PSQI was found for Night 2, (*r_s_*(43) = -.39, *p* = .008, *q* = .025), and the average between Night 1 and Night 2, (*r_s_*(43) = -.41, *p* = .005, *q* = .021). For semantically unrelated morning recall scores a significant correlation was found for Night 1, (*r_s_*(46) = -.40, *p* = .005, *q* = .021), Night 2 (*r_s_*(44) = -.31, *p* = .037, *q* = .075), and the average between nights (*r_s_*(44) = -.41, *p* = .005, *q* = .021), however, Night 2 morning recall for unrelated pairs did not survive correction for multiple comparisons. For semantically related pre-sleep recall scores, Night 2, (*r_s_*(42) = -.38, *p* = .010, *q* = .051), and the average were significant, (*r_s_*(41) = -.30, *p* = .050, *q* = .118) but Night 1 was not, (*r_s_*(46) = -.17, *p* = .244, *q* = .244). However, none of these results survived correction. For unrelated pre-sleep recall scores, Night 1, (*r_s_*(49) = -.38, *p* = .007, *q* = .039), and the average were significant, (*r_s_*(45) = -.34, *p* = .021, *q* = .085), but Night 2 was not, (*r_s_*(45) = -.28, *p* = .059, *q* = .118). Only Night 1 scores for unrelated word-pairs remained significant after correction. No other comparisons approached significance for semantically related or unrelated morning recall scores, *p* > .05. No scores for daily morning sleep quality measures were associated with either semantically related or unrelated word-pairs, *p* > .05.

For both semantically related and unrelated word-pairs, no correlations with AMI approached significance for any measure. Similarly, no daily morning motivation measures were associated with either semantically related or unrelated scores, *p* > .05.

## DISCUSSION

The present study examined the stability of overnight memory consolidation across multiple nights using a novel smartphone application. Contrary to our hypothesis, overnight retention did not demonstrate significant intra-individual stability across nights, despite significant consistency in pre-sleep learning, pre-sleep recall, and morning recall performance. Overnight retention was similarly unstable across nights for both semantically related and unrelated word pairs, while pre-sleep recall and morning recall showed significant intra-individual consistency across nights for unrelated word pairs only. Pre-sleep learning rate exhibited significant intra-individual consistency across nights for both related and unrelated word pairs. Furthermore, even after averaging scores across nights to reduce night-to-night variability, sleep quality was not associated with overnight retention, however, was associated with morning recall scores. PSQI scores were also associated with performance for semantically related and unrelated word pairs.

In contrast to the significant across-night consistency observed for pre-sleep learning, pre-sleep recall, and morning recall, overnight retention showed considerably greater night-to-night variability. The lack of significant correlation and a weak ICC of .35 suggests that consolidation performance was not a stable within-person characteristic across nights. This was corroborated by Bland-Altman analysis, which indicated that while there was little systematic change across nights, the wide limits of agreement highlight that there were large within-person fluctuations between nights. This finding contrasts with previous research suggesting that cognitive abilities show substantial trait-like stability (Breit et al., 2024). Both pre-sleep learning and recall measures demonstrated no mean-level differences and significant correlations between nights, in line with previous test-retest reliability literature of same-day memory tasks such as the RAVLT and VPA, which involve similar learning and recall components (Lee et al., 2023; Magalhaes et al., 2012). However, it is important to note that these studies only tested the stability of learning and recall scores, not overnight retention measures. The strong effect size and narrow limits of agreement of pre-sleep learning rate and recall suggest that this stability in scores is likely a reflection of trait-like encoding ability, consistent with literature showing that verbal learning slope is predictive of long-term recall (Walhovd et al., 2020). The observed night-to-night variability in overnight retention may therefore reflect both theoretical characteristics of the consolidation process and methodological features of its measurement.

Firstly, variability in SDMC may be the result of fluctuating daily factors. Transient factors such as stress may contribute to the findings in this sample, being particularly prevalent in university students (Knapp et al., 2024). Stress can negatively impact sleep quality in some individuals (Kalmbach et al., 2020). Sleep and cortisol interactions have also been highlighted to influence memory consolidation for information that has been attended to (Bennion et al., 2015). Additionally, night-to-night fluctuations in sleep quality may have influenced overnight retention. Although daily morning subjective sleep ratings were not associated with overnight retention measures, these measures were broad and were not available for the entire sample. Objective sleep quality, including sleep fragmentation or proportion of deep NREM sleep may have varied between nights. Future studies should include repeated measures of both objective sleep quality and SDMC to assess if sleep variability may influence SDMC. Finally, it is important to note that consolidation is not a process that occurs purely during sleep. Participants completed their learning and pre-sleep recall tasks approximately 3 hours before their bedtime, leaving a substantial interval for activities post-recall. A meta-analysis by Weng et al. (2025) highlighted that wakeful rest after learning benefits memory consolidation compared to engagement in other activities. Thus, participants may have had demanding post-recall tasks one evening (e.g. studying) and rested the next, possibly contributing to the lack of stability in our findings.

From a methodological perspective, the apparent instability of SDMC may partly reflect how overnight retention was quantified. Overnight retention was calculated as a percentage of morning recall relative to pre-sleep recall, meaning that the resulting ratio incorporates variability from both measurements and may be particularly sensitive when pre-sleep recall is relatively low. These considerations are particularly pertinent when considering that the word-pair task used in the present study was adapted from Lam et al. (2021), who utilised 32 word-pairs in adults with mild cognitive impairment. In contrast, many studies examining SDMC have employed tasks with upwards of 80 word-pairs (Denis et al., 2023; Ong et al., 2020; Wilhelm et al., 2020). For paradigms with fewer word-pairs, each additional item recalled or forgotten represents a relatively larger change in percentage performance; consequently, small differences in recall between nights may translate into comparatively large differences in overnight retention. Some of the observed night-to-night variability in overnight retention may therefore reflect limited measurement resolution and the operationalisation of the overnight retention score itself, rather than variability in consolidation alone. Consistent with this, morning recall demonstrated less variability across nights than pre-sleep recall. This suggests that instability in overnight retention was driven by variability in the denominator as opposed to variability that emerges following the overnight retention interval. Notably, both pre-sleep and morning recall remained more stable across nights than the derived SDMC score, suggesting that instability may be particularly associated with quantifying change between the two time points. To ensure observed overnight retention instability was not a result of test timing differences between nights, timing checks were conducted across nights. No significant difference in timing was found for pre-sleep learning, pre-sleep recall, or morning recall start times. Despite this, the gap between learning and pre-sleep recall times significantly differed across nights. Reassuringly, neither the difference in learning and pre-sleep recall times or the difference in morning recall times were correlated with overnight retention or morning recall, suggesting that instability is unlikely due to test start times.

Interestingly, the stability of both pre-sleep recall and morning recall varied according to semantic relatedness. For unrelated word pairs, pre-sleep and morning recall were significantly correlated across nights and demonstrated good ICCs. In contrast, neither pre-sleep nor morning recall for related word pairs were stable between nights, despite showing narrower limits of agreement. Despite appearing counterintuitive, this dissociation may be partly explained by ceiling effects. As the word lists used were originally designed for older populations with mild cognitive impairment (Lam et al., 2021), semantically related pairs may have been insufficiently challenging in this sample. The apparently greater stability of unrelated word-pair performance should therefore be interpreted cautiously, as it may reflect the greater range and discriminatory capacity of the more difficult condition rather than an inherent difference in the stability of memory for unrelated versus semantically related information.

These findings are important to consider in the context of experimental studies designed to manipulate sleep in order to influence SDMC. Some sleep-manipulation studies have reported no significant differences between manipulation and control conditions (Bueno-Lopez et al., 2019; Diep et al., 2020; Eggert et al., 2013; Harrington et al., 2021; Henin et al., 2019; Koo et al., 2018; Omlin et al., 2018; Paßmann et al., 2016; Sahlem et al., 2015; Schneider et al., 2020; van Sluijs et al., 2020; Wilhelm et al., 2020), while others have demonstrated improved word-pair consolidation following interventions targeting sleep-dependent processes (Marshall et al., 2006; Ngo et al., 2013; Schreiner & Rasch, 2015; Papalambros et al., 2017; Perrault et al., 2019; Westerberg et al., 2015). It is important to note that a meta-analysis by Harlow et al. (2023) found a yearly decline in effect sizes for manipulation studies, suggesting earlier positive findings may have been a result of publication bias or changing methodology. Although null effects have sometimes been interpreted as suggesting that SDMC is relatively resistant to external manipulation, the present findings highlight an alternative consideration: substantial endogenous night-to-night variability may make experimentally induced changes more difficult to detect. If an individual’s overnight retention performance fluctuates considerably even in the absence of an intervention, differences observed between experimental and control nights may reflect both the effect of the manipulation and naturally occurring variation in consolidation. Conversely, a genuine manipulation effect may be obscured by this background variability, particularly in studies relying on a single experimental and control night. The mixed findings across manipulation studies may therefore reflect not only differences in the effectiveness of the interventions themselves, but also variability in the underlying consolidation process. Establishing an individual’s typical range and stability of SDMC across repeated baseline nights may consequently be important for distinguishing experimentally induced changes from normal night-to-night fluctuations.

The hypothesis that sleep quality would be associated with more robust overnight retention measures averaged across multiple measurements was partially supported. Average performance across nights (normalised overnight retention score) was not associated with sleep quality measured by the PSQI. However, the strength of this non-significant association remained consistent across nights, and there was no association between PSQI scores and variability in overnight retention. The lack of association is not entirely unexpected and may likely be attributed to the nature of the PSQI measure. The PSQI is a self-report index that captures overall sleep quality in the past month. As consolidation processes have been posited to rely on neural activity during sleep, such as spindles, oscillations, and synaptic downscaling (Born & Wilhelm, 2012; Tononi & Cirelli, 2014), the broad construct measured by the PSQI may not align with the specific processes involved in SDMC each night. Indeed, subjective sleep is not reliably predictive of objective sleep (Cudney et al., 2022), and the two measures of sleep have been found to diverge in their ability to predict memory (Cavuoto et al., 2016), potentially explaining the lack of association. It should be noted that morning recall scores for both mornings and their average were associated with PSQI and remained consistent with moderate effect sizes, such that poorer sleep quality resulted in poorer recall. Similarly, averaged pre-sleep recall scores were associated with PSQI. Supporting this, subjective sleep quality has previously been shown to influence academic and cognitive performance, including attention and executive function (Adelantado-Renau et al., 2019; Ampofo et al., 2025). As a result, PSQI may reflect broader, trait-like cognitive processes that are captured in recall scores but not in overnight retention scores. For both semantically related and unrelated word pairs, PSQI was significantly correlated with the averaged morning recall. This suggests that the association between subjective sleep quality and SDMC may not be dependent on the strength of memories learned. Thus, the findings for overnight retention scores and raw scores suggest that while subjective sleep quality may influence recall, it does not reliably predict SDMC. Future studies may benefit from investigating the influence of objective sleep quality on SDMC across multiple nights using PSG or actigraphy, to determine if specific sleep measures predict performance.

Despite the novel contributions of the study to SDMC research, there are several limitations that should be acknowledged in the context of these findings. Firstly, the word pair task used for the study was adapted from Lam et al. (2021), who designed the task for older adults with mild cognitive impairment. While the task is suitable for older adults, it may have increased the sensitivity to detect smaller fluctuations in SDMC variability. Future studies assessing stability in young healthy populations should consider incorporating a learning criterion and longer lists to better capture variability in SDMC. Furthermore, while the study increased ecological validity by administering tasks through a novel smartphone application, this methodology also comes with limitations, mainly concerning experimental control. Different environments in which participants completed the learning and recall tasks, may have contributed confounding factors of performance such as noise (Thompson et al., 2022) or environmental distractions that may have impacted attention and results across nights (Stothart et al., 2015). These distractions may have also contributed to the incompletion of tasks that would otherwise be completed if in a lab setting, thus affecting pairwise analyses. Future studies should aim to control for these confounders for the duration of tasks. Additionally, the present study did not control for whether participants utilised any encoding strategies for word-pairs. Strategies such as integrative encoding (i.e. imagining word-pairs in a scene) have been shown to greatly benefit subsequent retention after sleep (Denis et al., 2023). Future studies should ask participants if such strategies were used, allowing for more nuanced analyses. Finally, there were no objective assessments of sleep quality. While our analyses attempted to assess the influence of nightly sleep quality on SDMC performance, these were done through simple subjective morning ratings, limiting power and missing objective information such as nightly awakenings. Future studies should couple objective measures (e.g. wearable devices) with word-pair testing over multiple nights to assess the influence of objective sleep characteristics on SDMC performance.

## Conclusion

To our knowledge, this is the first study to examine the stability of SDMC over multiple nights. Notably, our findings highlight that the SDMC process may exhibit greater variability than previously assumed, subsequently calling into question the reliance on one-night SDMC protocols. Studies investigating SDMC should consider retesting over multiple nights to ensure robust findings. Clinically, these findings highlight the importance of repeated assessments in the diagnosis of memory consolidation issues. Our findings emphasise the need for future studies to further examine modifiable factors that underlie SDMC variability across nights, particularly in older adults, as these may be targeted for interventions.

## Acknowledgements

We thank all the participants from the Healthy Brain Ageing Clinic who contributed their time to being a part of this study. We acknowledge the contribution of all the clinical and administrative staff members at the Healthy Brain Ageing Clinic, the Woolcock Institute of Medical Research, and the Brain and Mind Centre.

The data that supports the findings of this study are openly available in the Sydney eScholarship Repository at https://doi.org/10.25910/3y8s-yg14.

For the data analysis stage of this manuscript, ChatGPT-4/5 was used to help revise and debug R code. All revised and debugged code was thoroughly tested and verified by authors.

